# MHC class IIa haplotypes derived by high throughput SNP screening in an isolated sheep population

**DOI:** 10.1101/2020.07.20.212225

**Authors:** K.L. Dicks, J.M. Pemberton, K.T. Ballingall, S.E. Johnston

**Author notes:** Corresponding author Kara Dicks, Royal Zoological Society of Edinburgh, 134 Corstorphine Road, Edinburgh, EH12 6TS. Present address: Royal Zoological Society of Scotland, Edinburgh, EH12 6TS, UK.

## Abstract

Investigating the current evolutionary processes acting on a highly polymorphic gene region, such as the major histocompatibility complex (MHC), requires extensive population data for both genotypes and phenotypes. The MHC consists of several tightly linked loci with both allelic and gene content variation, making it challenging to genotype. Eight class IIa haplotypes have previously been identified in the Soay sheep (*Ovis aries*) of St. Kilda using Sanger sequencing and cloning, but no single locus is representative of all haplotypes. Here, we exploit the closed nature of the island population of Soay sheep and its limited haplotypic variation to identify a panel of SNPs that enable imputation of MHC haplotypes. We compared MHC class IIa haplotypes determined by Sanger sequence-based genotyping of 135 individuals to their SNP profiles generated using the Ovine Infinium HD BeadChip. A panel of 11 SNPs could reliably determine MHC diplotypes, and two additional SNPs within the *DQA1* gene enabled detection of a recombinant haplotype affecting only the SNPs downstream of the expressed genes. The panel of 13 SNPs was genotyped in 5951 Soay sheep, of which 5349 passed quality control. Using the Soay sheep pedigree, we were able to trace the origin and inheritance of the recombinant SNP haplotype. This SNP-based method has enabled the rapid generation of locus-specific MHC genotypes for large numbers of Soay sheep. This volume of high-quality genotypes in a well-characterized population of free-living sheep will be valuable for investigating the mechanisms maintaining diversity at the MHC.

## INTRODUCTION

The most highly polymorphic region of the vertebrate genome is the major histocompatibility complex (MHC), and yet the evolutionary processes driving this diversity remain much debated (Bernatchez and Landry 2003; Spurgin and Richardson 2010). MHC genes encode proteins that present pathogen-derived peptides to T-cells, thus enabling an individual to mount an immune response (Klein 1986). Pathogen-mediated selection (PMS) is strongly implicated in driving diversity at the MHC (Bernatchez and Landry 2003; Piertney and Oliver 2006; Spurgin and Richardson 2010). The key PMS mechanisms involved, which are not necessarily mutually exclusive, are heterozygote advantage (Doherty and Zinkernagel 1975; Slade and McCallum 1992) and variation in selection pressure through either negative frequency-dependent selection (Apanius et al. 1997) or fluctuating selection (Hill 1991; Hedrick 2002). Disentangling these mechanisms is challenging in experimental systems which may be incapable of replicating the wide array of pathogens and parasites that occur within a wild host (although see Kalbe et al. 2009 and Eizaguirre et al. 2012), and within wild systems which may be swamped with variation in individual ontogeny, environment, genetic background and pathogen diversity.

A major impediment to understanding current selection patterns on the MHC in natural populations is our inability to accurately characterize variation at the level of the individual, and to do this at scale across many individuals. Developing genotyping methods which attribute alleles to loci and loci to haplotypes are key technical challenges when studying the MHC in non-model species (Bernatchez and Landry 2003; Spurgin and Richardson 2010). Large sample sizes are necessary to generate sufficient power in statistical analyses of highly polymorphic loci, such as the MHC. Rare alleles, which are of particular interest in studies of the MHC as they may confer increased fitness under negative frequency-dependent selection or in heterozygote genotypes under heterozygote advantage (Apanius et al. 1997; Hill 1998), enhance the need for large sample sizes. Characterization of each locus individually is required to partition variation amongst loci and determine an individual’s true genotype. Using pooled diversity measures generated by co-amplification of multiple indistinguishable loci prohibits assessment of the heterozygous state of each locus, and therefore the ability to effectively assess heterozygous advantage (Spurgin and Richardson 2010). Locus-based genotyping of the MHC, particularly at the haplotype level, is often expensive, time consuming or not possible due to gene duplication, gene conversion and variation in gene number which makes designing locus-specific primers difficult or impossible (Westerdahl 2007; Babik 2010; Whittaker et al. 2012, but see Worley et al. 2008). Moreover, MHC alleles exist in haplotypes and, in the short-term, selection will act on the combination of alleles present in haplotypes or diplotypes. Genotyping only a single locus may not capture the full diversity of the MHC molecules generated by single haplotypes (e.g. Dicks et al. 2019). An effective and reliable genotyping method should, therefore, be able to capture variation at individual loci and across multiple expressed loci.

The Soay sheep (*Ovis aries*) population of St. Kilda, UK provides an excellent study system in which to look for signatures of the evolutionary mechanisms operating to maintain diversity at the MHC. Soay sheep are descendants of an early domestication and have been unmanaged on the island of Soay for probably thousands of years, and since the translocation of 107 sheep to the adjacent island of Hirta in 1932 (Clutton-Brock et al. 2004). The population has been the subject of a long-term study since 1985 (Clutton-Brock et al. 2004). Multiple population censuses per year and the sampling of lambs in spring and of sheep of all age categories in summer has provided a wealth of data on both phenotypic and fitness traits relevant to understanding signatures of evolutionary processes acting on the MHC, as well as a deep understanding of their ecology and demography. Driven primarily by climate and food limitation at high density, the population undergoes large fluctuations in population density (Boyd et al. 1964), varying between approximately 400 and 2000 sheep on the island of Hirta. However, the effective population size was estimated at 194 sheep (Kijas et al. 2012) and the population has generally low genetic diversity compared to other breeds (Lawson Handley et al. 2007; Kijas et al. 2009). This intensively studied island population which is subjected almost exclusively to natural evolutionary processes lends itself to investigating MHC evolution. Indeed, a previous study found that allele frequencies of two MHC-linked microsatellite loci were more even that would be expected under neutral processes, implicating balancing selection (Paterson 1998), and associations have been detected between the MHC-linked microsatellites and parasite resistance (fecal egg counts) and fitness traits (survival in lambs and yearlings) (Paterson et al. 1998).

Sequence-based genotyping methods have been developed for multiple ovine MHC class IIa loci (Ballingall and Tassi 2010; Ballingall et al. 2015, 2018), and variation at class IIa loci was previously characterized in the Soay sheep population using these methods (Dicks et al. 2019). Genotyping of the MHC class II-*DRB1*, *DQA1, 2* and *2-like*, as well as *DQB1, 2* and *2-like* loci revealed eight haplotypes, with some allele sharing amongst haplotypes (but not between loci; see Figure S1). No single locus captures all MHC class IIa variation in the Soay sheep population. In addition, sequence-based genotyping of the *DQ* loci, especially the *DQB* loci, was complicated by cross-amplification amongst loci and variation in locus configuration (Ballingall et al. 2015, 2018; Ali et al. 2016; Dicks et al. 2019). Sanger sequencing of multiple loci in large numbers of individuals would be costly and slow, yet such haplotypic data is necessary to carry out analyses of the evolutionary processes underlying the high variability in this region.

SNP-based genotyping methods are easily scalable, and human population studies have used selected SNPs which are in linkage disequilibrium with specific MHC haplotypes to impute MHC haplotypes (Dilthey et al. 2011; de Bakker and Raychaudhuri 2012; Zheng et al. 2014). High linkage disequilibrium between loci within subregions is characteristic of the mammalian MHC (Dawkins et al. 1999), including the ovine MHC (Lee et al. 2012). However, high levels of polymorphism and sequence similarity between alleles at different loci within the class IIa genes limits the ability to select SNPs from within coding regions, and sequence data for introns and other non-coding regions of the ovine class IIa is generally lacking. As a consequence, the Illumina Ovine 50K SNP BeadChip has just one SNP in the putative MHC class IIa region and the Illumina Ovine Infinium HD BeadChip, with an attempted 606,066 SNPs, has just 27 (Figure 1).

**Figure 1.**
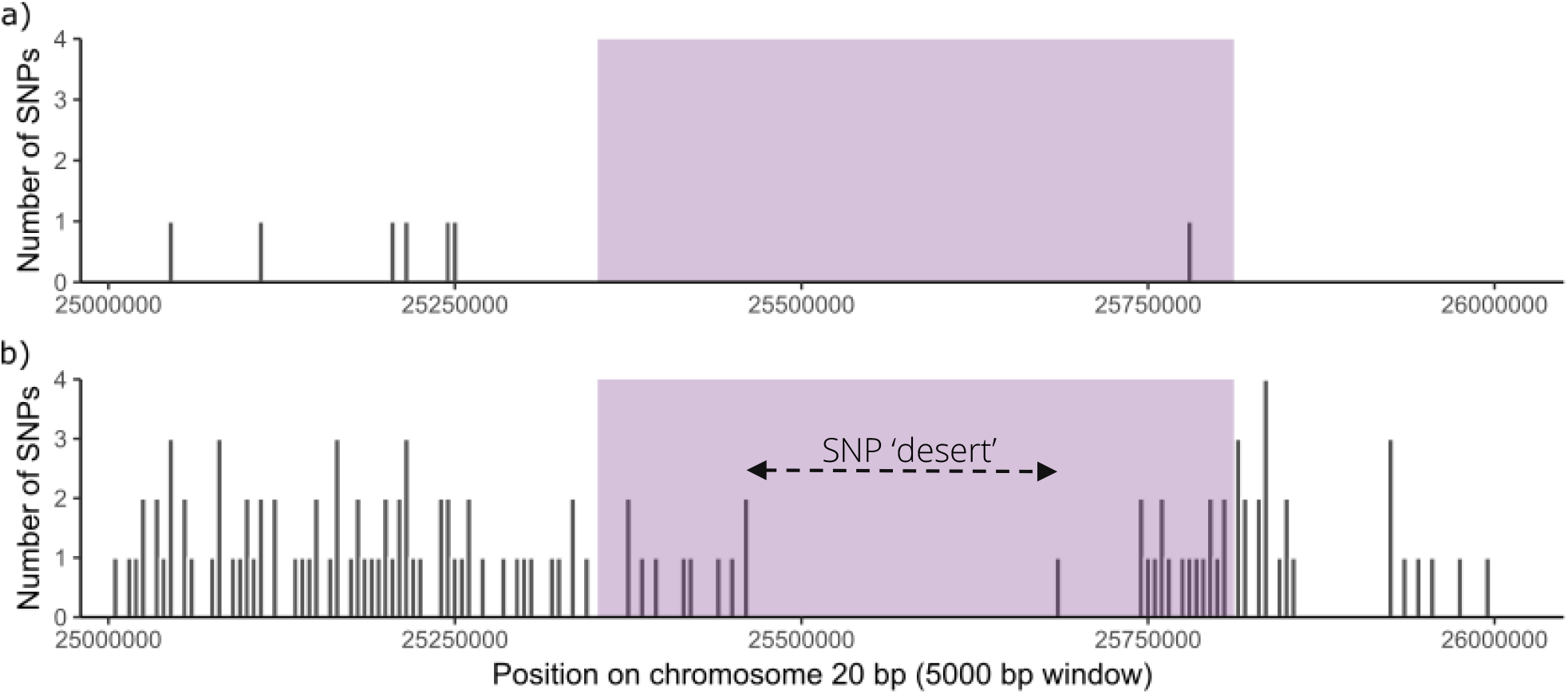
Number of SNPs within each 5000 bp window on chromosome 20 between 25 and 26 Mb on a) the 50K SNP chip and b) the HD SNP chip. The purple block represents the class IIa gene region between 25.353 to 25.812 Mb (determined here – see results). A SNP ‘desert’, an extended region lacking SNPs on the HD SNP chip, is shown by the dashed arrow.

Here, we develop a high throughput haplotyping system for the Soay sheep MHC class IIa. We hypothesized that on the Illumina Ovine Infinium HD BeadChip there might be a sufficient number of SNPs in the flanking regions that were in linkage disequilibrium with the class IIa region to enable imputation of the Soay haplotypes. We then aimed to identify a panel of SNPs capable of imputing the Soay sheep haplotypes, and to genotype them in a large number of individuals spanning the 30-year study period for which DNA was available.

## MATERIALS AND METHODS

### Study system and DNA sampling

The Soay sheep living in the Village Bay are of Hirta, St. Kilda, UK have been intensively monitored since 1985 (Clutton-Brock et al. 2004). Lambs within the study area of Village Bay are caught, ear-tagged and sampled for genetic analysis soon after birth. Individuals which move into the study area as adults are sampled when possible, during spring, August catch-up or the autumn rut. Individual life-histories are determined through lambing observations, censuses of the Village Bay population during spring, August and autumn, and regular mortality checks. The pedigree of Soay sheep has previously been defined using 315 SNPs selected from the 50K SNP chip combined with mother-offspring relationships observed during lambing (Bérénos et al. 2014).

### Ovine Infinium HD SNP BeadChip and MHC class IIa genotyping

Johnston et al. (2016) previously genotyped 188 sheep at 606 066 SNPs on the Ovine Infinium HD SNP BeadChip (2016). These sheep were selected from the pedigree in order to maximize the genetic diversity captured by the fewest individuals.

Soay sheep MHC class IIa haplotypes were characterized by Dicks et al. (2019) using Sanger sequence-based genotyping of individual loci. We sequenced the *DRB1* and *DQA1, DQA2* and DQA*2-like* loci in half (n=94) of the sheep that were also genotyped on the Ovine Infinium HD SNP BeadChip, and we sequenced *DRB1* and *DQA1* in the remaining half (n=94) individuals. *DQA2*, *DQA2-like* and *DQB* loci were not genotyped due technical challenges with co-amplification of multiple loci, so it was not possible to distinguish between haplotypes A and G.

### Identification of SNPs linked to MHC class IIa haplotypes

The exact locations of the MHC genes on chromosome 20 are poorly defined, so we first estimated the MHC class IIa gene region in the *O. aries* genome (Oar_v3.1, GCA_000298735.1) on Ensembl (Yates et al. 2016). We searched for protein families including the words “class II histocompatibility” in their description, limiting results to chromosome 20 and unmapped scaffolds, and using the single-copy gene *BTNL2* to define the q-arm telomeric end of the class IIa region (Gao et al. 2010; Liu et al. 2011; Lee et al. 2012). The Soay sheep Ovine Infinium HD SNP BeadChip genotypes were then filtered using PLINK v1.90 (Purcell et al. 2007) to select polymorphic SNPs within the identified MHC class IIa region with a genotyping rate > 95%, and a minor allele frequency < 1 %.

Pairwise linkage disequilibrium (LD) between SNPs within 500 kb of each other was calculated in Haploview (Barrett et al. 2005) as both *r*^2^ and *D*’. LD blocks were identified using the Four Gamete Rule (Wang et al. 2002). Blocks were numbered according to their proximity to the SNP ‘desert’ in the center of the class II region (Figure 1), with those upstream (i.e. closer to the centromere) being negative and those downstream (i.e. closer to the q-arm telomere) being positive. The seven SNPs within block −1 were phased using BEAGLE (Browning and Browning 2007) with standard settings and compared to the class IIa haplotypes. Upstream (more negative) SNPs were then added sequentially to block −1 SNP haplotypes and phased using BEAGLE after each SNP addition. This identified if and where the SNP and class IIa haplotypes matched, and where LD degraded, i.e. where a single class IIa haplotype was represented by multiple SNP haplotypes. The process was repeated for SNPs downstream of the SNP desert.

A recombination between haplotypes B and H was identified (see results), such that the SNP profile matched haplotype H but the class IIa haplotype was haplotype B. To enable identification of this recombinant haplotype using SNPs alone, the second exons of class IIa genes were searched to identify SNPs suitable for a genotyping assay – biallelic across all haplotypes with a minimum of 50 bp of flanking sequence in each direction containing limited variation.

### KASP SNP panel selection and genotyping

Candidate SNPs were validated *in silico* using Kraken software by LGC Genomics (Hoddesdon) for Kompetitive allele specific PCR (KASP). Of the approved SNPs, alleles and allelic combinations were identified which were unique to each class IIa Sanger haplotype, preferentially selecting those closest to the SNP ‘desert’ in the center of the class IIa region. A minimum subset of SNPs was identified which were able to impute the class IIa diplotype of an individual with degeneracy. The final panel of SNPs used for KASP genotyping included 11 SNPs selected from the Ovine Infinium HD BeadChip and two intragenic *DQA1* SNPs (Table 1).

**Table 1.** Locus information, genotyping success rate (%), MAF (minor allele frequency), HO (observed heterozygosity), HE (expected heterozygosity), and Hardy-Weinberg Equilibrium (HWE) significance values, for the SNPs selected for KASP genotyping. SNP statistics were calculated for the 5349 individuals which passed quality control. Genome positions for DQA1 loci are unknown.

| SNP ID | rs accession<sup>a</sup><br>or GenBank<br>accession<br>number | Genome<br>Position | Major<br>Allele | Minor<br>Allele | Genotyping<br>success<br>rate (%) | MAF | H<sub>O</sub> | H<sub>E</sub> | HWE<br>p-value |
| --- | --- | --- | --- | --- | --- | --- | --- | --- | --- |
| oar3_OAR20_25759002 | rs401547149 | 25759002 | G | A | 98.38 | 0.170 | 0.286 | 0.282 | 0.802 |
| oar3_OAR20_25458354 | rs403182119 | 25458354 | C | G | 99.38 | 0.123 | 0.205 | 0.215 | 0.327 |
| oar3_OAR20_25447962 | rs407228358 | 25447962 | A | G | 97.63 | 0.439 | 0.499 | 0.492 | 0.809 |
| oar3_OAR20_25393230 | rs408059130 | 25393230 | G | A | 99.25 | 0.145 | 0.254 | 0.248 | 0.636 |
| oar3_OAR20_25742711 | rs408458106 | 25742711 | G | A | 97.25 | 0.485 | 0.490 | 0.500 | 0.670 |
| oar3_OAR20_25441205 | rs412204478 | 25441205 | T | C | 98.38 | 0.473 | 0.517 | 0.499 | 0.476 |
| oar3_OAR20_25417300 | rs412489249 | 25417300 | G | A | 97.88 | 0.243 | 0.335 | 0.368 | 0.060 |
| oar3_OAR20_25761264 | rs414259673 | 25761264 | C | A | 98.25 | 0.442 | 0.502 | 0.493 | 0.738 |
| oar3_OAR20_25419505 | rs419818757 | 25419505 | T | C | 98.38 | 0.031 | 0.063 | 0.061 | 0.677 |
| oar3_OAR20_25742763 | rs421513037 | 25742763 | C | T | 97.75 | 0.490 | 0.494 | 0.500 | 0.813 |
| oar3_OAR20_25782804 | rs422511469 | 25782804 | G | A | 94.51 | 0.279 | 0.403 | 0.402 | 0.952 |
| DQA1_171 | LR025209 | 171<sup>b</sup> | C | G | 83.52<sup>c</sup> | 0.146 | 0.118 | 0.249 | <0.0001 |
| DQA1_195 | LR025209 | 195<sup>b</sup> | C | A | 82.77<sup>c</sup> | 0.148 | 0.116 | 0.252 | <0.0001 |

<sup>a</sup> Accession number in Ensembl http://www.ensembl.org/Ovis_aries/
<sup>b</sup> Position within GenBank sequence
<sup>c</sup> SNP carries known null-allele

DNA was extracted from tissues (ear punch or *post-mortem*) or buffy coat samples for a previous study using the Illumina Ovine50K SNP chip (Bérénos et al. 2014). The Qiagen DNeasy 96 Blood and Tissue Kit was used following the manufacturer’s protocol, with the exception that the final elution was performed using two 50 μL elutions (Bérénos et al. 2014). DNA was normalized to 50 ng/μL for 5951 individuals and distributed across 66, 96-well plates. A single sheep – the ‘golden sheep’ (ID 7568) – was included on 61 plates to confirm genotyping consistency and a no template control (NTC) was included on all plates. An additional 200 individuals were genotyped more than once for reasons unrelated to this work (for example when DNA was used for genotyping on the Ovine50K SNP chip, the DNA failed at the first attempt and so was re-attempted on a subsequent plate – note that differences in genotyping technologies between Illumina arrays and KASP may mean that a sample could be successful with one method but not the other). LGC Genomics performed KASP genotyping and automatic genotype calling for all 66 plates.

### Quality control and SNP haplotyping

Quality control was carried out excluding *DQA1* intragenic SNPs as they carry an expected null allele. PLINK v1.90 (Chang et al. 2015) was used to exclude individuals with more than 50% missing genotypes. Genotypes of the ‘golden sheep’ (ID 7568) were compared across all plates to identify any systematic errors. For other duplicated individuals, DNA was expected to be of poor quality and therefore more susceptible to allelic dropout, and so the genotyping attempt with the highest match to parental genotypes was included. For individuals genotyped in triplicate, the genotyping attempt with fewest mismatches to alternative attempts was retained. Assessments of the multiple genotyping attempts of the ‘golden sheep’ and 201 repeated individuals, as well as Mendelian error checks between offspring and parents indicated a high genotyping error rate on DNA plates 1-3, the very oldest plates, and so all genotypes from these plates were discounted.

Individuals with more than two (of 11) missing genotypes were excluded, and SNPs were phased using BEAGLE v4.0 (Browning and Browning 2007). Phased SNP haplotypes were assigned to their corresponding class IIa haplotypes. Twenty-six individuals with novel haplotypes were excluded as these were presumed genotyping or phasing errors (maximum novel haplotype frequency was four). A Mendelian inheritance check was carried out on the inferred haplotypes for each individual using the Soay sheep pedigree (Bérénos et al. 2014). Three individuals whose phased MHC diplotypes mismatched the diplotype of at least one parent were excluded from further analysis. 5349 sheep passed quality control measures.

### Detection of B-H recombinants

The class IIa gene *DQA1* is not present in all haplotypes (Ballingall et al. 2018; Dicks et al. 2019) and consequently the *DQA1* SNPs are effectively tri-allelic – they have a major allele, a minor allele, and a null allele where the DQA1 locus is absent from the haplotype. A *DQA1* genotype failure can, therefore, represent both true failures and the diplotype of an individual carrying two *DQA1*null* haplotypes. For all sheep identified as carrying haplotype H, the *DQA1* SNP genotypes were assessed to identify B–H recombinants.

### Hardy-Weinberg Equilibrium tests

Hardy-Weinberg equilibrium (HWE) predicts genotype (or diplotype) frequencies under random mating at loci with no selection, mutation or migration (Guo and Thompson 1992), and therefore, deviation from HWE would indicate that natural selection (including heterozygote advantage and negative frequency dependent selection) or sexual selection (including MHC based mate choice) may be acting on the haplotypes (Hedrick 2004), or that some other demographic process is acting (Guo and Thompson 1992). As selection can act more or less at different life history stages (LH stages), we assessed HWE for all known conceived sheep (live born, still born, and fetuses of mothers who died over winter), live born (survived to be tagged within approx. first week of life), survived to first August (approx. 4 months) and survived to second August (approx. 16 months). Individuals were known to be alive if they were observed in the summer census or August catch, or if they had a known birth year prior to the observation year and were observed alive or known to have died at a later date. HWE tests were performed in the R package GenePop v1.0.5 (Rousset 2008) as probability tests (the exact test of Haldane 1954; Guo & Thompson 1992; Weir 1996) where the null hypothesis is that the same allele frequencies are observed) and *U* tests for heterozygote excess (where the alternative hypothesis is heterozygote excess Raymond & Rousset 1995) for each LH stages (all cohorts combined) and for each cohort (LH stages) or year (standing population). Sequential Bonferroni correction (Holm 1979) was used to account for multiple testing.

Unless otherwise stated, all bioinformatic processes analyses were carried out in R version 3.3.3.

## RESULTS

### Identification of SNPs linked to MHC class IIa haplotypes

Full class IIa diplotypes were determined by sequence-based genotyping for 135 out of the 188 individuals genotyped on the Ovine Infinium HD BeadChip. Partial diplotype information was determined for an additional 50 individuals which carried haplotypes A or G, but these haplotypes were not differentiated (using only *DRB1* and *DQA1* loci). Three individuals failed sequence-based genotyping.

Nine protein families represented by 22 transcripts on the Oar_v3.1 genome were identified with the term “class II histocompatibility” in their descriptions, as well as *BTNL2* (Table S1). Five transcripts were located on unmapped scaffolds. The remaining 17 transcripts were clustered in two regions of chromosome 20, 7.164 – 7.426 Mb and 25.353 – 25.812 Mb, with the former including primarily class IIb gene annotations (*DO* and *DM* genes) and the latter class IIa gene annotations (*DR* and *DQ* genes). *DQA* gene annotations were found in both class II regions. The unusual ruminant class IIa gene *DYA* is located in the former region, 7.164 – 7.426 Mb (Ballingall et al. 2001) and so it is likely that this gene has been poorly annotated. Therefore, it is more likely that the *DQA* gene is located in the latter class IIa region, 25.353 – 25.812 Mb, along with the *DR* and other *DQ* genes. Previous assemblies of this region using BAC cloning did not identify *DQ* loci outside of the class IIa region (Liu et al. 2006; Gao et al. 2010). The classical class IIa genes are therefore expected to be located between 25.353 and 25.812 Mb.

On the Ovine Infinium HD BeadChip, there were 11,757 HD SNPs located on chromosome 20, of which 8,420 SNPs passed quality control in the Soay sheep sample. Twenty-one SNPs were located in the class IIa region predicted above (Figure 1b).

Pairwise LD estimates of *D*’ for the SNPs between 25.353 and 25.812 Mb were typically very high (Figure 2). There was no obvious pattern to the degradation of LD, which would be suggestive of a recombination hotspot. On the other hand, *r*^*2*^ estimates, were typically very low (Figure S2) and were also uninformative in identifying linkage regions. Given the lack of obvious LD pattern, linkage blocks were identified using the Four Gamete Rule in Haploview (shown in Figure 2).

**Figure 2.**
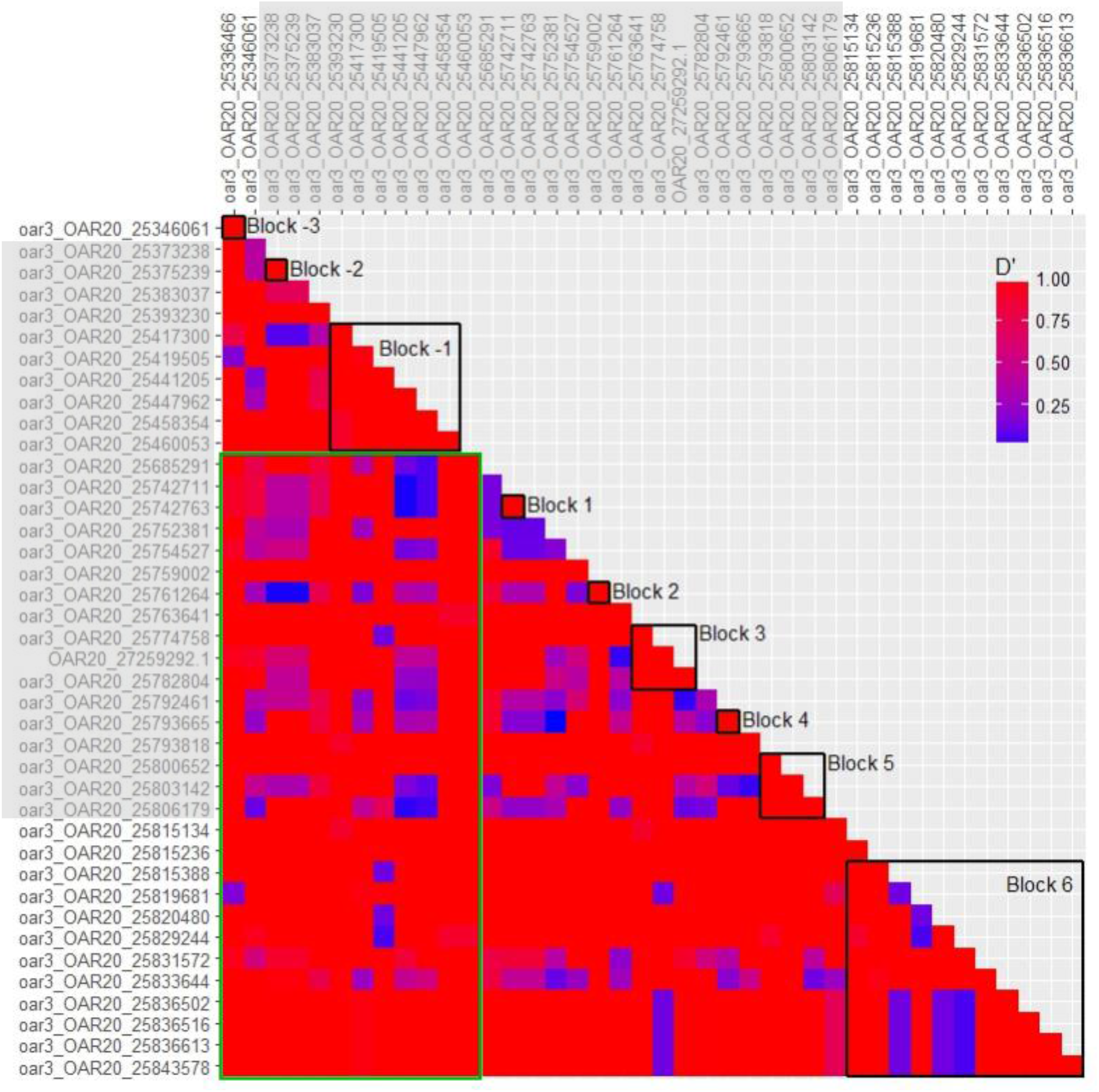
D’ linkage disequilibrium estimates for pairs of HD SNPs within and surrounding the class IIa gene region, where values of 1 represents high LD, and 0 represents low LD. SNPs within the class IIa gene region are highlighted by the grey shading. The SNP ‘desert’ is denoted by the green box, and SNP pairs that fall within this green box represent pairs spanning this SNP ‘desert’ (i.e. that one SNP is upstream and the other downstream). Open black boxes show linkage blocks identified within Haploview using the Four Gamete Rule.

Block –1, which is immediately upstream of the HD SNP gap, consisted of six SNPs and formed six haplotypes (Figure 3). Four of the six haplotypes in block –1 each corresponded to a single Sanger haplotype, and two each corresponded to a pair of haplotypes, A and G, or B and H (Figure 3). The addition of upstream SNPs to block –1 showed that linkage degraded immediately upstream of block –1 (at SNP oar3_OAR20_25383037) within haplotype E, but haplotypes A and G, as well as haplotypes B and H, only became differentiated at oar3_OAR20_25346061 (block ‒3). Downstream of the SNP ‘desert’, haplotypes B and H became differentiated at oar3_OAR20_25742763 (block 1), and A and G differentiated with the addition of oar3_OAR20_25752381 (adjacent to block 1). Linkage within haplotypes did not degrade until oar3_OAR20_25833644 (within block 6) for both haplotypes B and E (Figure 3). Thus, a continuous linkage region of 31 SNPs was identified between oar3_OAR20_25393230 and oar3_OAR20_25831572, which encompasses 438.34 kb.

**Figure 3.**
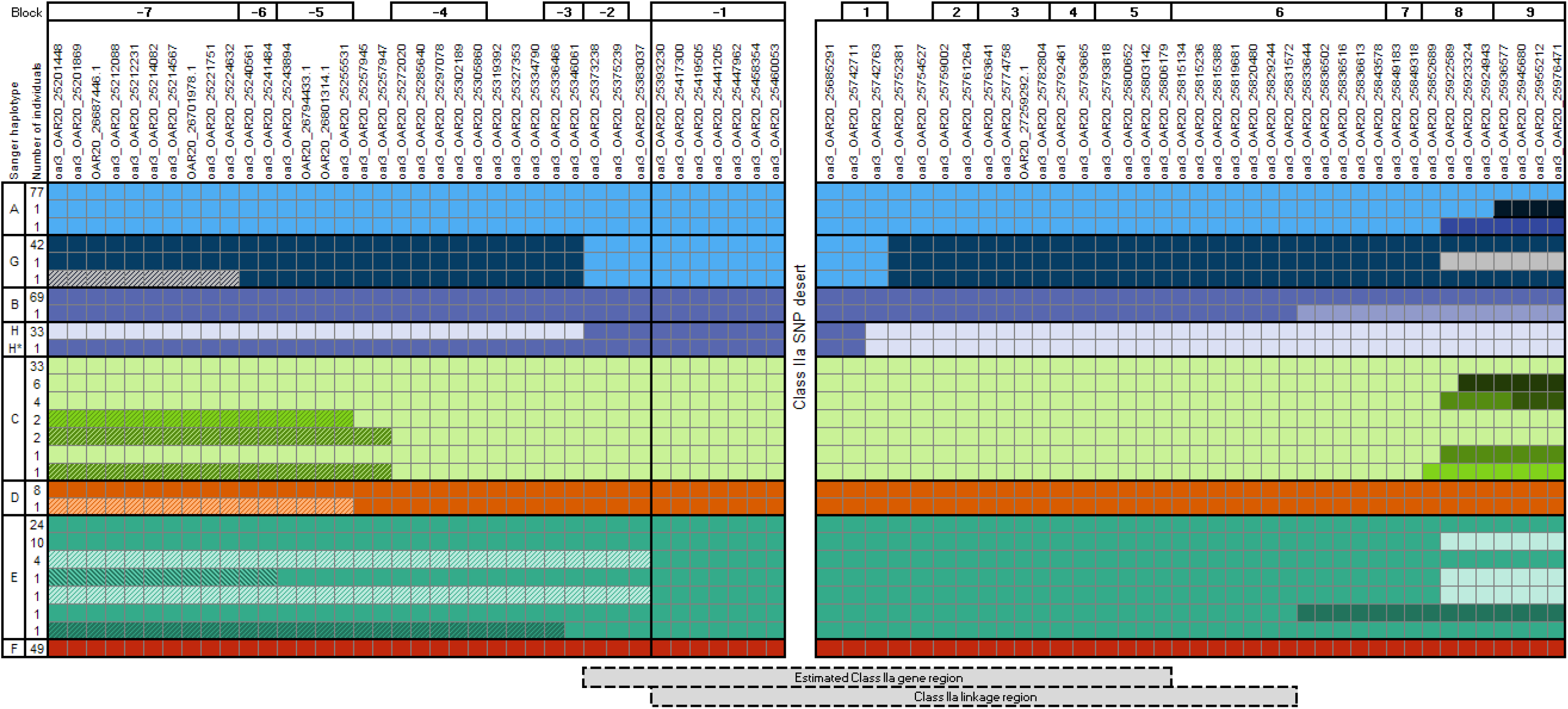
Representation of the phased SNP haplotypes (rows) formed by sequentially and cumulatively adding either upstream (left) or downstream (right) HD SNPs to block –1. Linkage blocks (Four Gamete Rule in Haploview) are indicated above the SNP names, and corresponding Sanger haplotypes in the left-most column. Moving outwards from block –1, a color change along the row indicates that an additional haplotype is formed with the addition of that SNP. The estimated class IIa gene region (25.353 to 25.812 Mb) and the class IIa linkage region are indicated below the plot. H* is a recombinant haplotype detected in a single individual, for which the SNP haplotype within the class IIa linkage region matches haplotype H but class IIa Sanger sequencing is haplotype B. Data are 188 sheep genotyped on the HD SNP chip and with MHC haplotypes established by sequence-based genotyping.

### Haplotype B and H recombinant detection

Following Sanger sequencing individual 4179 was found to be heterozygous for haplotypes for B and G. However, the flanking SNPs revealed a recombination event which did not affect the expressed haplotype but affected the haplotype that would be imputed using the panel of 31 SNPs within the class IIa linkage region. Using only the seven SNPs upstream of the SNP ‘desert’, 4179’s SNP diplotype was consistent with haplotypes B/H (identical in this region) and G (Figure 3, haplotype H*). Using only the 24 SNPs downstream of the SNP gap, 4179’s SNP diplotype was consistent with haplotypes H and G. Therefore, using the SNPs within the class IIa linkage region (i.e. block –1 and downstream SNPs), the haplotype would be incorrectly imputed as H. This indicates the existence of a recombination event between haplotypes B and H which has occurred downstream of the class IIa genes included in the Sanger haplotype.

### KASP assay selection and genotyping

All 31 SNPs within the class IIa linkage region that were selected from the Ovine Infinium HD SNP BeadChip passed *in silico* validation for KASP. From this shortlist, we selected 11 SNPs enabling imputation of the eight class II haplotypes whilst maintaining degeneracy in case of genotyping failures (Figure 4; Table 1). To enable detection of the B-H recombinant haplotype, two SNPs within exon 2 of the *DQA1* gene were identified (DQA1-171 and DQA1-195; Figure S3) that met the requirements for KASP assay design. No suitable biallelic SNPs with sufficient invariant flanking sequence could be identified in other MHC class IIa genes to make this distinction. *DQA1* is not present on three Soay sheep MHC class IIa haplotypes, A, E and G, which instead carry *DQA2-like* alleles. As such, *DQA1* SNPs are effectively tri-allelic and consist of a major allele, minor allele, and null allele (see Figure 4). 5951 individuals were genotyped by LGC Genomics using KASP at the final panel of 13 SNPs.

**Figure 4.**
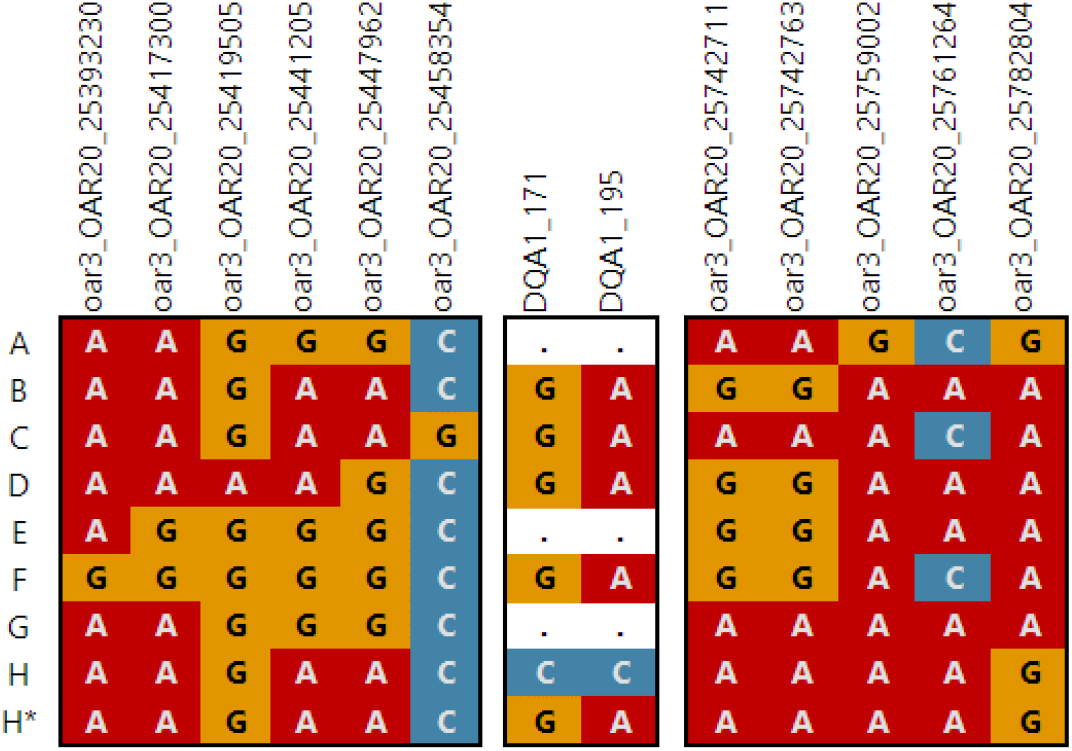
Alleles at KASP-genotyped SNPs (columns) within the MHC class IIa linkage region for each Sanger haplotype (rows). For DQA1 SNPs, dots correspond to DQA1*null alleles on haplotypes where the DQA1 locus is absent. DQA1*null alleles are therefore the absence of amplification. SNP loci are shown in order of their position on chromosome 20.

### Quality control

The ‘golden sheep’ was KASP genotyped 57 times, with an overall genotyping error rate of 0.48% and a maximum error rate of 1.85% per locus. After excluding DNA plates 1-3 due to exceptionally high error rates, 5675 individuals were KASP genotyped for the panel of 13 SNPs. Quality control measures on the 11 flanking SNPs, and samples sizes at each stage are shown in Figure S4. The genotyping rate was below 50% for 195 sheep, which were excluded.

After assessing genotyping attempts for repeated individuals and excluding the least successful genotyping attempt, 102 individuals were removed due to having four or more missing genotypes. The 11 flanking SNPs were phased in the remaining 5378 individuals. This revealed eight frequent haplotypes which exactly matched the eight expected class IIa haplotypes. Twenty novel haplotypes were identified in 26 individuals (with a maximum frequency of four, see Table S2) and were excluded from further analysis as they were considered probably due to genotyping errors or extremely rare haplotypes. Mendelian error checks identified only three sheep whose haplotypes mismatched their parents’; the offspring were removed from the dataset. The final dataset consisted of diplotypes for 5349 individuals. In this sample, all SNPs were in HWE, except for the *DQA1* SNPs which were expected to deviate from HWE due to null alleles (Table 1).

### *DQA1* SNP genotypes

*DQA1* SNP genotypes were compared to the diplotypes identified by the panel of 11 flanking SNPs. The two selected *DQA1* SNPs, *DQA1*-171 and *DQA1*-195, are tri-allelic such that haplotype H carries the minor allele (C for both loci), four haplotypes (B, C, D, and F) have the major allele (G or A respectively), and three haplotypes (A, E and G) do not carry the locus and present as null alleles. As such, it is possible to predict the expected genotypes for the *DQA1* SNPs from an individual’s diplotype. In a test sample of 801 sheep, five had unexpected *DQA1* genotypes, all of which carried *DQA1*\*null haplotypes (see Figure S5), which is an error rate of 0.62%. Additionally, genotypes classified as fail, which include true negatives, true failures to amplify, and homozygote null-alleles, showed greater variance in fluorescence for the *DQA1* SNPs compared to the panel of 11 truly bi-allelic SNPs. This suggests non-specific amplification may be occurring in the absence of the target locus. Nevertheless, B–H recombinant haplotypes could be identified (see Figure S5).

In the full dataset, genotypes for the two *DQA1* SNPs were assessed only in individuals carrying haplotype H (N=906), identifying 56 recombinant individuals. After recoding recombinant haplotypes as R, all recombinant individuals were found to be descendants of the individual in which the recombinant haplotype was identified, male 4179 (Figure 5). Furthermore, the mother of 4179 carried both B and H haplotypes, and thus was the likely origin of the cross-over event.

**Figure 5.**
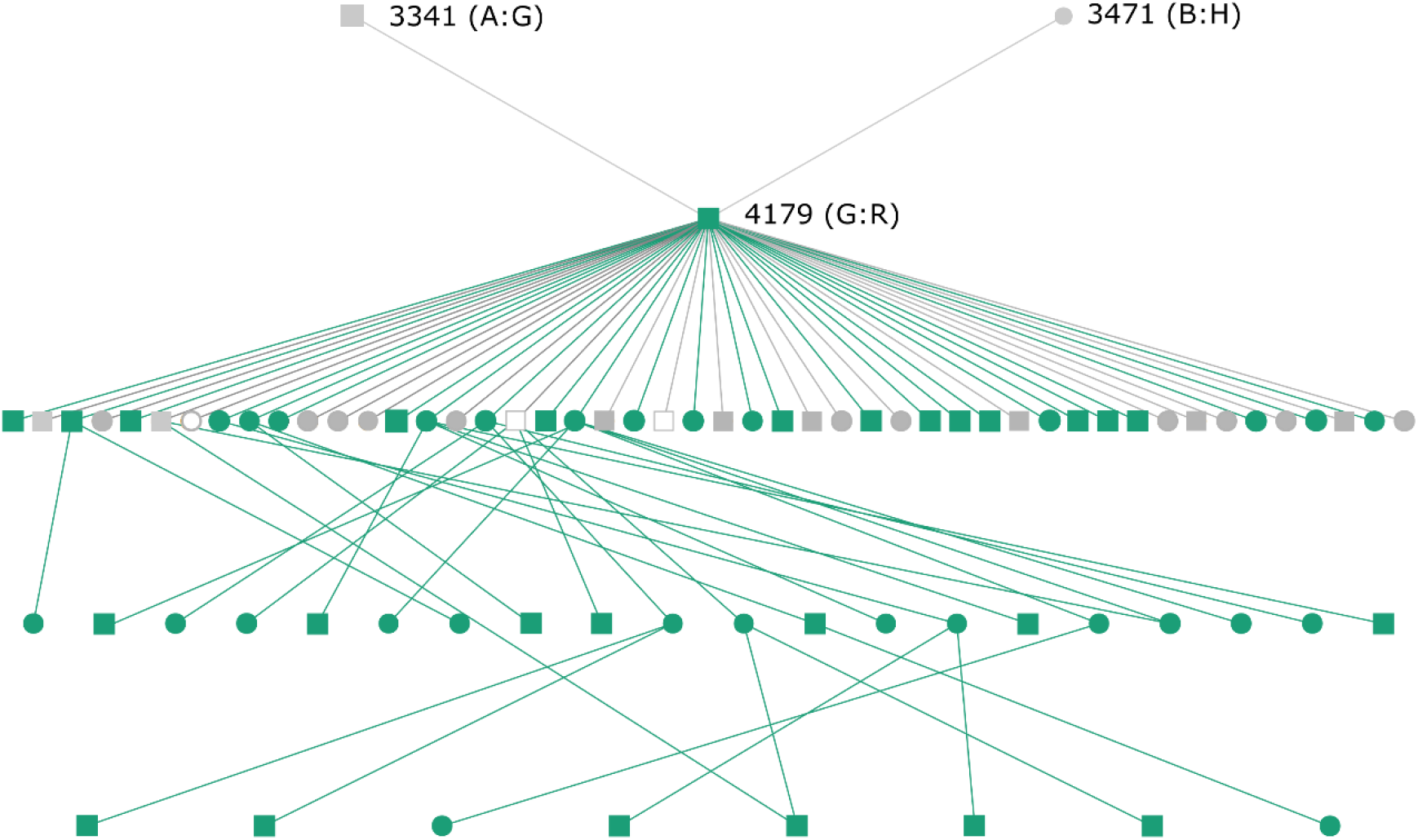
Pedigree of individual 4179, who carries the first identified B–H recombinant haplotype. Green represents individuals carrying the recombinant haplotype, grey not carrying the recombinant haplotype, and open represents individuals that have not been genotyped. Squares are males, circles are females. All offspring of ID 4179 are shown, but subsequent descendants are only shown if they carry the recombinant haplotype. Diplotypes of 4179 and his parents are shown in brackets.

### Haplotype and diplotype frequencies

Of 5349 individuals, 976 (17.9%) had homozygous diplotypes. Haplotype B was the most frequent, whilst haplotype D the rarest, with only six individuals homozygous for haplotype D (Figure 6b). All possible diplotypes were observed (Figure 6a). Haplotype frequencies for individuals born in 3-year periods from 1985 to 2012 (5137 individuals with known birth years) varied over time (Figure 7).

**Figure 6.**
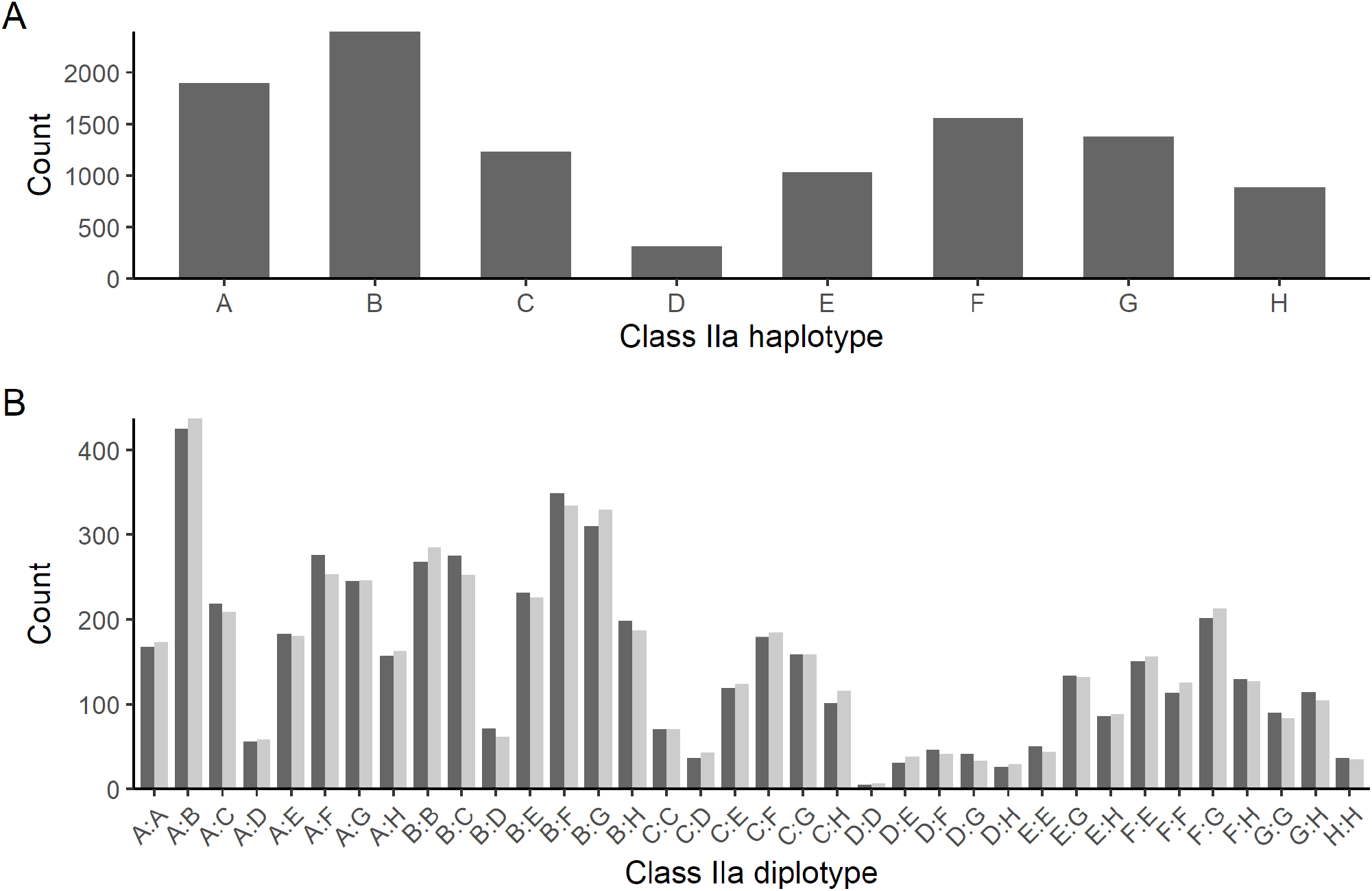
Observed haplotype frequencies (A), and observed (dark) and expected (light) diplotype frequencies under HWE (B). N= 5349 individuals in both cases.

**Figure 7.**
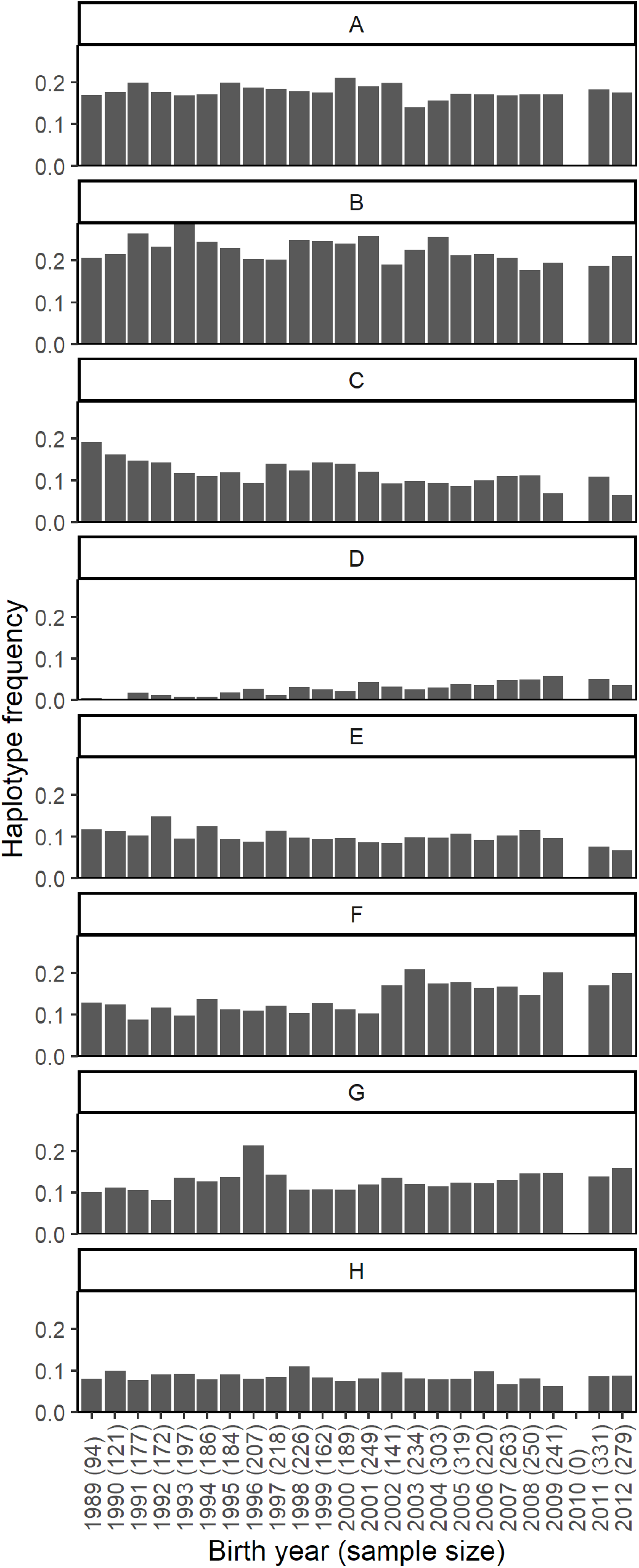
Haplotype frequencies in individuals born each year between 1989 and 2012. Sample sizes (number of individuals with MHC genotypes, i.e. half the number of haplotypes) are shown in brackets beside the birth year. Technical failure resulted in very few samples being haplotyped for individuals born in 2010, therefore frequency estimates are not shown.

### Hardy-Weinberg Equilibrium tests

None of the four life history stages considered were found to deviate from HWE (Table 2). HWE tests were carried out for each cohort independently at each of the four life stages. The 2012 ‘known conceived’ cohort was marginally out of HWE for the exact test (*p* = 0.023, α threshold = 0.025), with more homozygotes than expected (H_E_ = 235.30, H_O_ = 223), as was the 2004 ‘live born’ cohort (*p* = 0.021, α threshold = 0.025), also with more homozygotes than expected (H_E_ = 232.02, H_O_ = 241). All other years and tests for each LH did not deviate from HWE (Table S3).

**Table 2.** Results from HWE tests (exact and heterozygote excess – p values) for each LH stage. Sample sizes (number of individuals with MHC genotypes) are shown (n).

| LH stage | n | HWE |  |
| --- | --- | --- | --- |
|  |  | Exact test | Heterozygote excess |
| Known conceived | 5158 | 0.8275 | 0.7654 |
| Live born | 4643 | 0.7738 | 0.7462 |
| Survived to 1<sup>st</sup> August | 3872 | 0.6856 | 0.7193 |
| Survived to 2<sup>nd</sup> August | 1824 | 0.8801 | 0.4257 |

## DISCUSSION

Analyses of the evolutionary processes underlying variation at the MHC require population-scale data. To achieve this, a SNP-based genotyping method was developed for rapid and accurate determination of the class IIa diplotypes in Soay sheep. Using SNPs selected from the Ovine Infinium HD BeadChip, a linkage region surrounding the class IIa genes was identified, and from it a minimal panel of 11 SNPs was selected that would enable imputation of the class IIa haplotypes. Furthermore, a recombination event between haplotypes B and H was detected, and two intragenic SNPs were selected to detect such recombinant haplotypes. Finally, this panel of 13 SNPs was applied to 5349 Soay sheep using KASP genotyping technology.

The classical MHC class IIa genes were estimated as occurring between 25.353 and 25.812 Mb on the *O. aries* genome, although poor mapping within this region prevents identification of the exact positions of MHC genes. Nevertheless, twenty-one SNPs on the Ovine Infinium BeadChip were located within the identified class IIa gene region, with a 0.225 Mb ‘SNP desert’ located in the center of this region. This lack of SNPs is not a reflection of a lack of variable SNPs in this region but rather their unsuitability for incorporation on a SNP chip due to the presence of low minor allele frequencies, and variable flanking sequences that prevent the design of robust primers and probes.

Pairwise measures of LD, *D*’ and *r*^*2*^, showed opposing patterns across the MHC class IIa region. Pairwise *D*’ was typically very high, whilst pairwise *r*^*2*^ was typically very low in this region. This opposing pattern of *D*’ and *r*^*2*^ measures of LD was also found in the class IIa, IIb and III regions in Rylington Merino sheep (Lee et al. 2012). VanLiere and Rosenberg (2008) showed that *r*^*2*^ and *D*’ are not always directly related, and r^2^ is sensitive to MAF, especially MAF < 0.3 which is not uncommon within the MHC. Lee et al. (2012) showed that pairwise LD using *D*’ in Rylington Merino sheep was high within class IIa but decreased between the class IIa and class IIb regions, as well as between class IIa and class III regions. This suggests recombination within the class IIa is reduced compared to recombination between it and the neighboring MHC subregions. Pairwise LD measures were therefore perhaps unlikely to reveal a relationship between the class IIa haplotypes and flanking SNPs on the Ovine Infinium HD BeadChip.

An alternative method of assessing the pattern of LD between the flanking SNPs and the class IIa region, the Four Gamete Rule, was used to identify linkage blocks as a starting point to cumulatively add SNPs to determine where SNP haplotypes degraded. This method detected a linkage region of 31 SNPs that matched the class IIa haplotypes. This class IIa linkage region extended between 25.39 – 25.83 Mb according to the allocated physical positions of the SNPs, which sits well within the estimated class IIa gene region of 25.35 – 25.81 Mb. The downstream extent of the estimated class IIa gene region is marked by the gene *BTNL2*, which, because it is a single copy gene, is more likely to be accurately mapped than the other multi-copy genes in this region. The overlap between the estimated class IIa gene region and the SNP linkage region suggests that at least 24 of the 31 SNPs are located within the estimated class IIa region.

Diplotypes for the 5349 Soay sheep which passed quality control can be considered highly accurate. Indirect MHC typing methods, such as SNP genotyping, are faster and cheaper than direct sequencing methods but may incur an increased error rate which must be minimized. Phasing SNPs to determine the two SNP haplotypes carried by an individual, carried out here with BEAGLE (Browning and Browning 2007), uses information from all individuals within the dataset to calculate the most likely haplotypes. BEAGLE also uses this information to impute missing SNP genotypes. Therefore, phasing errors are increased when there is a high genotyping error rate, and thus errors can be reduced by using a more stringent genotyping rate cut-off across all SNPs. The repetition of a single individual on almost all DNA plates was valuable in detecting systematic errors on three plates, probably due to the DNA having been exhausted or degraded by freeze-thawing and revealed a genotyping error rate of 0.48% (after excluding plates 1–3). The Mendelian inheritance check demonstrated that the phased haplotypes were consistent between parents and offspring, with only three exceptions. Additionally, only 20 novel haplotypes were identified in 26 individuals (0.5 % of haplotyped individuals), which were probably due to genotyping errors, including allelic dropout. Whilst this suggests that genotyping errors can lead to incorrect haplotype inferences, in this population, with good pedigree information, it is clearly possible to detect such errors.

A recombination event between haplotypes B and H (Sanger haplotype B upstream, SNP haplotype H downstream) was identified in one individual (ID 4179) genotyped on the Ovine Infinium HD BeadChip. The recombination event must have occurred downstream of the expressed class IIa genes characterized within the Sanger haplotypes (*DRB1, DQA* and *DQB*) but upstream of the SNP oar3_OAR20_25742763, leaving the functional class IIa haplotype unaffected. Individuals were selected for genotyping on the Ovine Infinium HD BeadChip because they were highly representative of the genetic variation within the Soay sheep population (Johnston et al. 2016), and male 4179 was included because he was very prolific with a large number of descendants. Using two SNPs located within the exons of the class IIa locus *DQA1*, this recombination was detected in 54 of 4179’s descendants and, because both of 4179’s parents were genotyped, we were able to detect the origin of the recombination event during gametogenesis in his mother. The use of SNPs located within loci that are not present on every haplotype, as for the two *DQA1* loci, was less than ideal due to the increased error rate caused by non-specific amplification. Nevertheless, their inclusion on the KASP panel enabled the identification of a substantial proportion of recombinant individuals with high accuracy, as apparent by a lack of parent-offspring mismatches following recombinant calling. Were this recombination event not identified, and therefore not detectable, a substantial number of haplotypes would have been incorrectly imputed, which could have impacted subsequent analyses.

Whilst the *DQA1* SNPs were effective for detecting the identified downstream B to H recombination, it is unlikely to be possible to detect novel recombination events involving upstream SNPs between haplotypes A and G. The six SNPs located upstream of the class IIa genes and the *DQA1* SNPs were identical for these haplotypes (see **Figure 3** and **Figure 4**). All other haplotype pairs have unique upstream and downstream SNP profiles.

All possible diplotypes were identified within the population, although the rarest, homozygote for D, was only identified in six out of the 5349 individuals genotyped (0.11 %). Over the study period (1985 – 2012), 5137 haplotyped individuals had known birth years (or expected birth years in the case of fetuses). There was no evidence of dramatic shifts in haplotype frequencies, but they were found to vary over time. There may be a slight decrease in haplotype C and a slight increase in haplotype D (**Figure 7**) which warrants further investigation to identify whether these shifts are significant, and if so, if they are due to natural or selective forces.

There was no evidence of deviation from HWE for any of the LH stages assessed. Whilst HWE was also assessed in each cohort at each LH stage, sample sizes were often relatively small, which may limit the power to detect any deviations. Deviations from HWE were detected using the exact test for the ‘known conceived’ stage in 2012 and for the heterozygote excess test in at the ‘live born’ stage in 2004 (both an excess of homozygotes); however, significance was marginal after applying sequential Bonferroni correction and so some caution should be applied in considering this evidence of deviation from HWE. Whilst deviation from HWE can indicate selective or demographic processes operating within the population (Guo and Thompson 1992; Spurgin and Richardson 2010), lack of deviation from HWE does not necessarily provide good evidence to the contrary (Spurgin and Richardson 2010).

Genotyping the MHC is notoriously challenging (Bernatchez and Landry 2003; Piertney and Oliver 2006; Babik 2010; Spurgin and Richardson 2010) but accurate, locus-specific genotypes are vital for unravelling the modes of evolution operating to maintain diversity at the MHC. Previous work identified eight MHC class II haplotypes in the island population of Soay sheep (Dicks et al. 2019) using sequence-based genotyping, and here we have described a SNP-based (KASP) genotyping method which enabled rapid genotyping of 5349 sheep (out of a total 5675 sheep attempted). A panel of SNPs has the advantage of being relatively easy to genotype, suitable for high-throughput genotyping and does not suffer from artefacts, but it is susceptible to incorrect haplotype phasing due to genotyping errors, particularly allelic dropout which may be most prevalent in poor quality or quantity DNA (Gill 2001; Butler 2005). The stringent quality control measures applied here should minimize any effects of genotyping and phasing errors, and the Soay sheep pedigree was particularly valuable here in enabling the inclusion of a Mendelian inheritance check as a quality control measure. As a method for genotyping MHC in a non-model organism, however, this panel of SNPs required high initial optimization effort in terms of preliminary class IIa sequencing and is not easily transferable to other breeds of sheep or other species without extensive preliminary sequencing of haplotypes. Nevertheless, this method has provided a valuable haplotype-level MHC data set that is accompanied by extensive phenotypic and fitness data that will enable future studies on the evolutionary processes acting to maintain diversity at the MHC in a natural environment.

## ACKNOWLEDGEMENTS

We thank the National Trust for Scotland and the Scottish Natural Heritage for permission to work on St. Kilda and QinetiQ for logistics and support. Many thanks to Jill Pilkington and all Soay sheep project members and volunteers for collection of data and samples. Thanks to Phil Ellis and Camillo Bérénos who prepared DNA samples for Ovine SNP50 and HD BeadChip genotyping. KLD was supported by a Biotechnology and Biological Sciences Research Council (BBSRC) doctoral training grant (grant number BB/J01446X/1). KTB receives funding from the European Union’s Horizon 2020 research and innovation programme under grant agreement no. 731014KB (VetBioNet) and acknowledges the support received from the Scottish Government’s strategic research programme. The Soay Sheep Project is funded by the UK Natural Environment Research Council and the SNP chip genotyping was done at the Wellcome Trust Clinical Research Facility Genetics Core and funded by the European Research Council. There are no conflicts of interest.

## REFERENCES

Ali AOA, Stear A, Fairlie-Clarke K, et al (2016) The genetic architecture of the MHC class II region in British Texel sheep. Immunogenetics 1–7. doi: 10.1007/s00251-016-0962-6

Apanius V, Penn D, Slev PR, et al (1997) The nature of selection on the major histocompatibility complex. Crit Rev Immunol 17:179–224. doi: 10.1615/CritRevImmunol.v17.i2.40

Babik W (2010) Methods for MHC genotyping in non-model vertebrates. Mol Ecol Resour 10:237–251. doi: 10.1111/j.1755-0998.2009.02788.x

Ballingall KT, Lantier I, Todd H, et al (2018) Structural and functional diversity arising from intra and inter haplotype combinations of duplicated DQA and B loci within the Ovine MHC. Immunogenetics 70:257–269. doi: 10.1007/s00251-017-1029-z

Ballingall KT, MacHugh ND, Taracha ELN, et al (2001) Transcription of the unique ruminant class II major histocompatibility complex-DYA and DIB genes in dendritic cells. Eur J Immunol 31:82–86. doi: 10.1002/1521-4141(200101)31:1<82::AID-IMMU82>3.0.CO;2-X

Ballingall KT, Steele P, Lantier I, et al (2015) An ancient interlocus recombination increases class II MHC DQA diversity in sheep and other Bovidae. Anim Genet 46:333–336. doi: 10.1111/age.12290

Ballingall KT, Tassi R (2010) Sequence-based genotyping of the sheep MHC class II DRB1 locus. Immunogenetics 62:31–39. doi: 10.1007/s00251-009-0410-y

Barrett JC, Fry B, Maller J, Daly MJ (2005) Haploview: Analysis and visualization of LD and haplotype maps. Bioinformatics 21:263–265. doi: 10.1093/bioinformatics/bth457

Bérénos C, Ellis PA, Pilkington JG, Pemberton JM (2014) Estimating quantitative genetic parameters in wild populations: A comparison of pedigree and genomic approaches. Mol Ecol 23:3434–3451. doi: 10.1111/mec.12827

Bernatchez L, Landry C (2003) MHC studies in nonmodel vertebrates: What have we learned about natural selection in 15 years? J Evol Biol 16:363–377. doi: 10.1046/j.1420-9101.2003.00531.x

Boyd JM, Doney JM, Gunn RG, Jewell PA (1964) The soay sheep of the island of Hirta, St. Kilda. A study of a feral population. Proc Zool Soc London 142:129–164. doi: 10.1111/j.1469-7998.1964.tb05159.x

Browning SR, Browning BL (2007) Rapid and Accurate Haplotype Phasing and Missing-Data Inference for Whole-Genome Association Studies By Use of Localized Haplotype Clustering. Am J Hum Genet 81:1084–1097. doi: 10.1086/521987

Butler JM (2005) Forensic DNA typing: biology, technology, and genetics of STR markers, 2nd edn. Elsevier, Burlington

Chang CC, Chow CC, Tellier LCAM, et al (2015) Second-generation PLINK: rising to the challenge of larger and richer datasets. Gigascience 4:7. doi: 10.1093/bioinformatics/btu495

Clutton-Brock TH, Pemberton JM, Coulson T (2004) The sheep of St Kilda. In: Clutton-Brock TH, Pemberton JM (eds) Soay Sheep: Dynamics and selection in an island population. Cambridge University Press, Cambridge, pp 17–50

Dawkins R, Leelayuwat C, Gaudieri S, et al (1999) Genomics of the major histocompatibility complex: haplotypes, duplication, retroviruses and disease. Immunol Rev 167:275–304. doi: 10.1111/j.1600-065X.1999.tb01399.x

de Bakker PIW, Raychaudhuri S (2012) Interrogating the major histocompatibility complex with high-throughput genomics. Hum Mol Genet 21:R29–R36. doi: 10.1093/hmg/dds384

Dicks KL, Pemberton JM, Ballingall KT (2019) Characterisation of major histocompatibility complex class IIa haplotypes in an island sheep population. Immunogenetics 1–11. doi: 10.1007/s00251-019-01109-w

Dilthey AT, Moutsianas L, Leslie S, McVean G (2011) HLA*IMP--an integrated framework for imputing classical HLA alleles from SNP genotypes. Bioinformatics 27:968–972. doi: 10.1093/bioinformatics/btr061

Doherty PC, Zinkernagel RM (1975) Enhanced immunological surveillance in mice heterozygous at the H-2 gene complex. Nature 256:50–52. doi: 10.1038/256050a0

Eizaguirre C, Lenz TL, Kalbe M, Milinski M (2012) Divergent selection on locally adapted major histocompatibility complex immune genes experimentally proven in the field. Ecol Lett 15:723–731. doi: 10.1111/j.1461-0248.2012.01791.x

Gao J, Liu K, Liu H, et al (2010) A complete DNA sequence map of the ovine major histocompatibility complex. BMC Genomics 11:466. doi: 10.1186/1471-2164-11-466

Gill P (2001) Application of low copy number DNA profiling. Croat Med J 42:229–232

Hedrick PW (2002) Pathogen resistance and genetic variation at MHC loci. Evolution (N Y) 56:1902–1908. doi: 10.1111/j.0014-3820.2002.tb00116.x

Hill AVS (1991) HLA associations with malaria in Africa: some implications for MHC evolution. In: Klein J, Klein D (eds) Molecular evolution of the major histocompatibility complex. Springer, Berlin, pp 403–419

Hill AVS (1998) The immunogenetics of human infectious diseases. Annu Rev Immunol 16:593–617

Johnston SE, Bérénos C, Slate J, Pemberton JM (2016) Conserved Genetic Architecture Underlying Individual Recombination Rate Variation in a Wild Population of Soay Sheep (Ovis aries). Genetics 203:

Kalbe M, Eizaguirre C, Dankert I, et al (2009) Lifetime reproductive success is maximized with optimal major histocompatibility complex diversity. Proc R Soc London B Biol Sci 276:. doi: 10.1098/rspb.2008.1466

Kijas JW, Lenstra JA, Hayes B, et al (2012) Genome-Wide Analysis of the World’s Sheep Breeds Reveals High Levels of Historic Mixture and Strong Recent Selection. PLoS Biol 10:e1001258. doi: 10.1371/journal.pbio.1001258

Kijas JW, Townley D, Dalrymple BP, et al (2009) A Genome Wide Survey of SNP Variation Reveals the Genetic Structure of Sheep Breeds. PLoS One 4:e4668. doi: 10.1371/journal.pone.0004668

Klein J (1986) The Natural History of the Major Histocompatability Complex. Wiley, New York, NY

Lawson Handley LJ, Byrne K, Santucci F, et al (2007) Genetic structure of European sheep breeds. Heredity (Edinb) 99:620–631. doi: 10.1038/sj.hdy.6801039

Lee CY, Qin J, Munyard KA, et al (2012) Conserved haplotype blocks within the sheep MHC and low SNP heterozygosity in the Class IIa subregion. Anim Genet 43:429–437. doi: 10.1111/j.1365-2052.2011.02268.x

Liu H, Liu K, Wang J, Ma RZ (2006) A BAC clone-based physical map of ovine major histocompatibility complex. Genomics 88:88–95. doi: 10.1016/j.ygeno.2006.02.006

Liu K, Zhang P, Gao J, et al (2011) Closing a gap in the physical map of the ovine major histocompatibility complex. Anim Genet 42:204–207. doi: 10.1111/j.1365-2052.2010.02083.x

Paterson S (1998) Evidence for balancing selection at the major histocompatibility complex in a free-living ruminant. J Hered 89:289–294. doi: 10.1093/jhered/89.4.289

Paterson S, Wilson K, Pemberton JM (1998) Major histocompatibility complex variation associated with juvenile survival and parasite resistance in a large unmanaged ungulate population. PNAS 95:3714–3719. doi: 10.1073/pnas.95.7.3714

Piertney SB, Oliver MK (2006) The evolutionary ecology of the major histocompatibility complex. Heredity (Edinb) 96:7–21. doi: 10.1038/sj.hdy.6800724

Purcell S, Neale B, Todd-Brown K, et al (2007) PLINK: a tool set for whole-genome association and population-based linkage analyses. Am J Hum Genet 81:559–575. doi: 10.1086/519795

Slade RW, McCallum HI (1992) Frequency-Dependent Selection at MHC Loci. Genetics 132:861–862

Spurgin LG, Richardson DS (2010) How pathogens drive genetic diversity: MHC, mechanisms and misunderstandings. Proc R Soc London B Biol Sci 277:979–988. doi: 10.1098/rspb.2009.2084

VanLiere JM, Rosenberg NA (2008) Mathematical properties of the measure of linkage disequilibrium. Theor Popul Biol 74:130–137. doi: 10.1016/j.tpb.2008.05.006

Wang N, Akey JM, Zhang K, et al (2002) Distribution of recombination crossovers and the origin of haplotype blocks: the interplay of population history, recombination, and mutation. Am J Hum Genet 1227–34. doi: 10.1086/344398

Westerdahl H (2007) Passerine MHC: genetic variation and disease resistance in the wild. J Ornithol 148:469. doi: 10.1007/s10336-007-0230-5

Whittaker DJ, Dapper AL, Peterson MP, et al (2012) Maintenance of MHC Class IIB diversity in a recently established songbird population. J Avian Biol 43:109–118. doi: 10.1111/j.1600-048X.2012.05504.x

Worley K, Gillingham MA., Jensen P, et al (2008) Single locus typing of MHC class I and class II B loci in a population of red jungle fowl. Immunogenetics 60:233–247. doi: 10.1007/s00251-008-0288-0

Yates A, Akanni W, Amode MR, et al (2016) Ensembl 2016. Nucleic Acids Res 44:D710–D716. doi: 10.1093/nar/gkv1157

Zheng X, Shen J, Cox C, et al (2014) HIBAG—HLA genotype imputation with attribute bagging. Pharmacogenomics J 14:192–200. doi: 10.1038/tpj.2013.18

