## Supplementary Information for "MHC class IIa haplotypes derived by high throughput SNP screening in an isolated sheep population"

Table S1. Locations of transcripts within protein families containing "class II histocompatibility" in the description occurring on chromosome 20 of the *Ovis aries* genome (Oar_v3.1, GCA_000298735.1) or on unmapped scaffolds.

| Protein Family ID | Description | Chromosome/ scaffold | Min position (bp) | Max position (bp) | Additional annotation | Protein ID |
|---|---|---|---|---|---|---|
| PTHR19944_SF45 | H2 Class II histocompatibility antigen E beta chain precursor | AMGL01121385.1 | 21 | 5,321 | | ENSOARP00000017463 |
| PTHR19944_SF52 | Class II histocompatibility antigen beta chain precursor | JH922951.1 | 480 | 710 | Human homologue DQB | ENSOARP00000000037 |
| PTHR19944_SF54 | HLA Class II histocompatibility antigen DQ alpha chain precursor | AMGL01119849.1 | 1,997 | 5,240 | DQ | ENSOARP00000003175 |
| PTHR19944_SF52 | Class II histocompatibility antigen beta chain precursor | JH922521.1 | 2,724 | 3,046 | Human homologue DQB | ENSOARP00000020450 |
| PTHR19944_SF45 | H2 Class II histocompatibility antigen E beta chain precursor | JH923254.1 | 3,431 | 4,459 | | ENSOARP00000002509 |
| PTHR19944_SF45 | H2 Class II histocompatibility antigen E beta chain precursor | 20 | 7,164,298 | 7,178,949 | | ENSOARP00000007745 |
| PTHR19944_SF55 | HLA class II histocompatibility antigen, DQ alpha 2 chain precursor | 20 | 7,198,200 | 7,208,505 | DQA | ENSOARP00000007864 |
| PTHR19944_SF55 | HLA class II histocompatibility antigen, DQ alpha 2 chain precursor | 20 | 7,198,200 | 7,204,636 | DQA | ENSOARP00000007865 |
| PTHR19944_SF52 | Class II histocompatibility antigen beta chain precursor | 20 | 7,221,436 | 7,228,230 | Human homologue DQB | ENSOARP00000007912 |
| PTHR19944_SF43 | Class II histocompatibility antigen DO beta chain precursor MHC class II antigen DOB | 20 | 7,234,829 | 7,248,527 | DOB | ENSOARP00000008004 |
| PTHR19944_SF51 | Class II histocompatibility antigen beta chain precursor | 20 | 7,346,814 | 7,357,333 | DMB | ENSOARP00000008482 |
| PTHR19944_SF13 | Class II histocompatibility antigen alpha chain precursor | 20 | 7,367,386 | 7,372,035 | DMA | ENSOARP00000008555 |
| PTHR19944_SF44 | HLA class II histocompatibility antigen DO alpha chain precursor MHC DN alpha MHC DZ alpha MHC class II antigen DOA | 20 | 7,420,413 | 7,425,930 | DOA | ENSOARP00000008678 |
| PTHR19944_SF54 | HLA Class II histocompatibility antigen DQ alpha chain precursor | 20 | 25,353,186 | 25,356,656 | DQ | ENSOARP00000016611 |
| PTHR19944_SF45 | H2 Class II histocompatibility antigen E beta chain precursor | 20 | 25,398,827 | 25,402,957 | | ENSOARP00000016782 |
| PTHR19944_SF45 | H2 Class II histocompatibility antigen E beta chain precursor | 20 | 25,450,028 | 25,460,293 | | ENSOARP00000016804 |
| PTHR19944_SF55 | HLA class II histocompatibility antigen, DQ alpha 2 chain precursor | 20 | 25,501,894 | 25,572,798 | DQA | ENSOARP00000016862, ENSOARP00000016864 |
| PTHR19944_SF52 | Class II histocompatibility antigen beta chain precursor | 20 | 25,532,506 | 25,543,103 | Human homologue DQB | ENSOARP00000017039 |
| PTHR19944_SF45 | H2 Class II histocompatibility antigen E beta chain precursor | 20 | 25,594,470 | 25,608,591 | DRB3 | ENSOARP00000017287 |
| PTHR19944_SF55 | HLA class II histocompatibility antigen, DQ alpha 2 chain precursor | 20 | 25,666,699 | 25,676,792 | DQA | ENSOARP00000017463 |
| PTHR19944_SF45 | H2 Class II histocompatibility antigen E beta chain precursor | 20 | 25,736,126 | 25,741,866 | | ENSOARP00000017715 |
| PTHR19944_SF47 | Class II histocompatibility antigen DR alpha chain precursor MHC class II antigen DRA | 20 | 25,774,090 | 25,783,626 | DRA | ENSOARP00000017840 |
| PTHR24100_SF5 | Butyrophilin like 2 | 20 | 25,793,691 | 25,811,932 | BTNL2 | ENSOARG00000016780 |

Table S2. Phased SNP haplotypes, their associated class IIa haplotype and the frequency of each haplotype. There were 20 novel SNP haplotypes at frequencies which did not match known class IIa haplotypes (denoted as NA). Lower case in the SNP profile for novel haplotypes indicates where the profile is one base different from a known class IIa haplotype.

| | SNP profile | Class IIa haplotype | Frequency |
|---|---|---|---|
| Known profiles | AACCGGATGCG | A | 1904 |
| | AACTAGGCAAA | B | 2337 |
| | AACTACATACA | C | 1233 |
| | AATTGGGCAAA | D | 317 |
| | AGCCGGGCAAA | E | 1041 |
| | GGCCGGGCACA | F | 1562 |
| | AACCGGATAAA | G | 1385 |
| | AACTAGATAAG | H | 949 |
| Novel profiles | AGCCGGGCAcA | NA | 4 |
| | AACCGGATAcA | NA | 3 |
| | AaCCGGGCACA | NA | 2 |
| | AACtGGATGCG | NA | 2 |
| | GGCCGGGCgCA | NA | 2 |
| | AACCGGgTGCG | NA | 1 |
| | AACTAGATAAa | NA | 1 |
| | AACTAGATAcA | NA | 1 |
| | AACTAGATgAG | NA | 1 |
| | AACTAGGCAcA | NA | 1 |
| | AACTAGGCgAA | NA | 1 |
| | AACtAGgTAAG | NA | 1 |
| | AACtGGATAAA | NA | 1 |
| | AAtCGGATAAA | NA | 1 |
| | AAtCGGATGCG | NA | 1 |
| | AGCCGGGCgAA | NA | 1 |
| | AgCTAGGCAAA | NA | 1 |
| | AgTCGGGCAAA | NA | 1 |
| | gACTAGGCAAA | NA | 1 |
| | GGtCGGGCACA | NA | 1 |

Table S3. Results from HWE tests (exact and heterozygote excess – p values) for each LH stage for each cohort. Sample sizes (number of individuals with MHC genotypes) are shown (n). Significant p values following sequential Bonferroni correction are highlighted in bold.

| | Known conceived | | | Live Born | | | Survived to 1st August | | | Survived to 2nd August | | |
|---|---|---|---|---|---|---|---|---|---|---|---|---|
| | | Exact test | Heterozygote excess | | Exact test | Heterozygote excess | | Exact test | Heterozygote excess | | Exact test | Heterozygote excess |
| Birth Year | n | P value | P value | n | P value | P value | n | P value | P value | n | P value | P value |
| 1989 | 94 | 0.1863 | 0.8455 | 94 | 0.0917 | 0.9088 | 68 | 0.0329 | 0.7467 | 51 | 0.1241 | 0.7553 |
| 1990 | 121 | 0.978 | 0.2055 | 121 | 0.9775 | 0.2041 | 116 | 0.9875 | 0.1478 | 100 | 0.9864 | 0.2284 |
| 1991 | 177 | 0.9477 | 0.0848 | 177 | 0.9301 | 0.1025 | 160 | 0.9136 | 0.207 | 42 | 0.3631 | 0.5565 |
| 1992 | 172 | 0.4599 | 0.181 | 172 | 0.5481 | 0.1988 | 97 | 0.1244 | 0.3845 | 75 | 0.0823 | 0.0832 |
| 1993 | 197 | 0.8644 | 0.64 | 197 | 0.872 | 0.6357 | 168 | 0.9257 | 0.5961 | 71 | 0.9794 | 0.2035 |
| 1994 | 186 | 0.6854 | 0.7723 | 186 | 0.7467 | 0.7897 | 133 | 0.7287 | 0.6986 | 23 | 0.3413 | 0.4374 |
| 1995 | 184 | 0.3743 | 0.0906 | 184 | 0.564 | 0.1261 | 137 | 0.8982 | 0.0649 | 120 | 0.8658 | 0.1276 |
| 1996 | 207 | 0.1353 | 0.8939 | 207 | 0.1957 | 0.8683 | 185 | 0.0958 | 0.9039 | 64 | 0.5661 | 0.3911 |
| 1997 | 218 | 0.4091 | 0.4227 | 218 | 0.4027 | 0.421 | 189 | 0.1665 | 0.1659 | 59 | 0.546 | 0.3651 |
| 1998 | 226 | 0.6183 | 0.9772 | 226 | 0.6257 | 0.975 | 202 | 0.2708 | 0.9932 | 14 | 0.2176 | 0.962 |
| 1999 | 162 | 0.7191 | 0.8833 | 162 | 0.764 | 0.8866 | 132 | 0.8812 | 0.9161 | 106 | 0.7093 | 0.9633 |
| 2000 | 189 | 0.2599 | 0.6619 | 189 | 0.2409 | 0.659 | 161 | 0.8449 | 0.5364 | 128 | 0.7988 | 0.4579 |
| 2001 | 249 | 0.7479 | 0.5833 | 249 | 0.6969 | 0.5221 | 245 | 0.6996 | 0.5316 | 15 | 0.0113 | 0.676 |
| 2002 | 141 | 0.1273 | 0.7734 | 141 | 0.1218 | 0.7791 | 131 | 0.1908 | 0.7603 | 117 | 0.2018 | 0.6822 |
| 2003 | 234 | 0.1714 | 0.9879 | 234 | 0.1802 | 0.9884 | 211 | 0.106 | 0.9951 | 171 | 0.1054 | 0.9446 |
| 2004 | 303 | 0.177 | 0.0265 | 303 | 0.1555 | **0.0206** | 210 | 0.0174 | 0.037 | 13 | 0.9731 | 0.3029 |
| 2005 | 319 | 0.8388 | 0.1847 | 319 | 0.5457 | 0.1891 | 135 | 0.7427 | 0.2111 | 62 | 0.947 | 0.0352 |
| 2006 | 220 | 0.0952 | 0.3741 | 220 | 0.0807 | 0.191 | 166 | 0.0962 | 0.2103 | 57 | 0.6511 | 0.1575 |
| 2007 | 263 | 0.5154 | 0.0236 | 263 | 0.3793 | 0.0626 | 153 | 0.1407 | 0.0305 | 85 | 0.0967 | 0.2232 |
| 2008 | 250 | 0.2011 | 0.1926 | 250 | 0.2596 | 0.2415 | 190 | 0.114 | 0.3734 | 91 | 0.2978 | 0.5806 |
| 2009 | 241 | 0.9721 | 0.2726 | 241 | 0.9894 | 0.2786 | 177 | 0.9668 | 0.1486 | 90 | 0.1886 | 0.1973 |
| 2011 | 331 | 0.2125 | 0.4636 | 331 | 0.4124 | 0.5648 | 229 | 0.5482 | 0.4589 | 14 | 0.8996 | 0.5606 |
| 2012 | 279 | **0.0232** | 0.9599 | 279 | 0.2627 | 0.8421 | 111 | 0.3796 | 0.8114 | 102 | 0.1864 | 0.8891 |

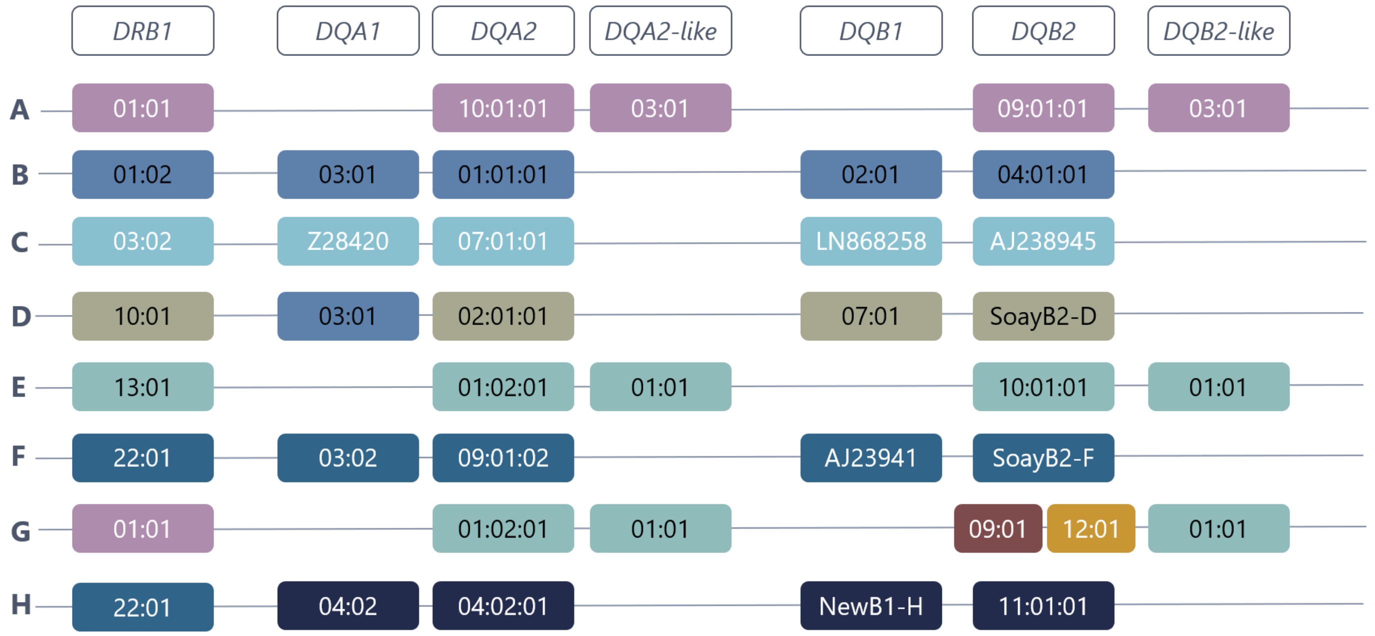


Figure S1. Visualization of the eight MHC class II haplotypes identified by (Dicks et al. 2019). Each row represents a haplotype and columns represent loci. Each allele is indicated according to its nomenclature (Ballingall et al. 2018; Dicks et al. 2019), which begins at 01:01 for each locus (i.e. *DRB1**01:01 is a different allele from *DQA2-like**01:01). Where the nomenclature is not available, the database accession number is provided. Colors are used solely to help visualize similarity where an allele at a locus is the same across haplotypes. Some alleles are shared across haplotypes (vertical), but not between loci (horizontal). Haplotypes either carry *DQA1* and *DQB1* or *DQA2-like* and *DQB2-like*; haplotypes carrying both loci have not been identified in Soay sheep. Haplotype G was found to have two *DQB2* alleles suggesting a gene duplication event; although one allele was identified from cDNA (RNA) and the other from genomic DNA and it is possible that only one is transcribed.

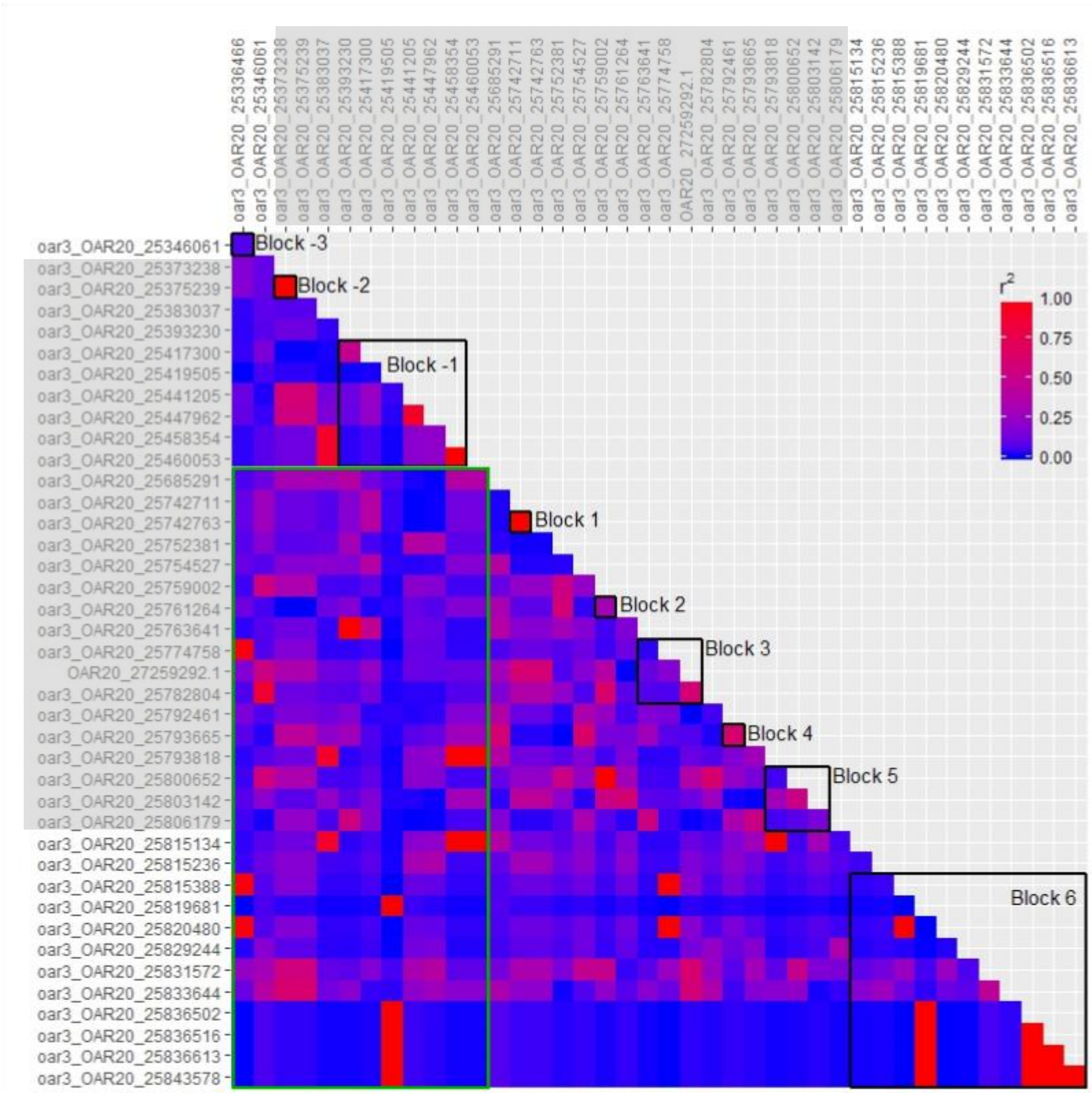


Figure S2. $r^2$ linkage disequilibrium estimates for pairs of HD SNPs within and surrounding the class IIa gene region, where values of 1 represents high LD, and 0 represents low LD. SNPs within the class IIa gene region are highlighted by the grey shading. The HD SNP gap is denoted by the green box, and SNP pairs that fall within this green box represent pairs spanning the SNP gap (i.e. that one SNP is upstream and the other downstream). Open black boxes show linkage blocks identified within Haploview using the Four Gamete Rule.

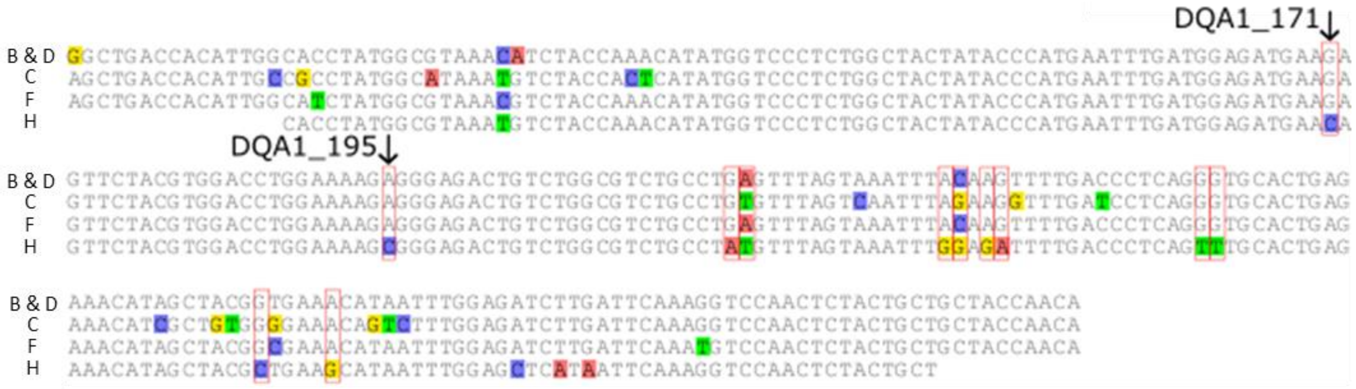


Figure S3. Aligned nucleotide sequences for exon 2 of *DQA1* alleles within the Soay sheep population for haplotypes which carry the *DQA1* locus. SNPs which distinguish haplotypes B and H are highlighted by red boxes. SNPs were only considered if they were bi-allelic across all alleles and at least 50 bp of flanking sequence on either side of the SNP was known. SNPs approved by KASP *in silico* trials are indicated by the arrows and SNP ID is shown beside the arrows.

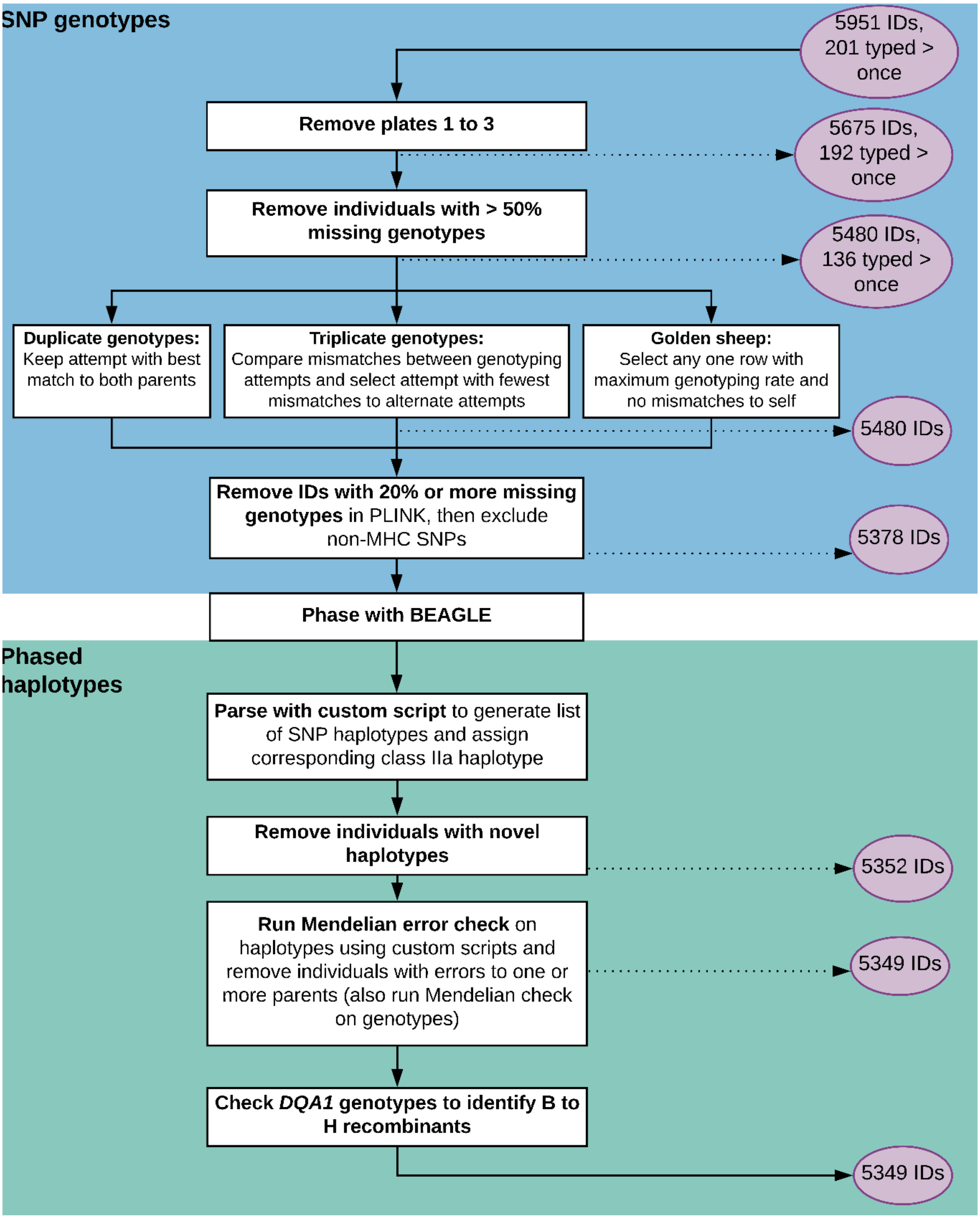


Figure S4. Flow chart illustrating the bioinformatic workflow for QC of KASP genotypes, showing processes in rectangles and the number of individuals at each stage in circles. QC processes using the SNP genotypes are shown in the top blue box, and using the phased haplotypes phased in the bottom green box.

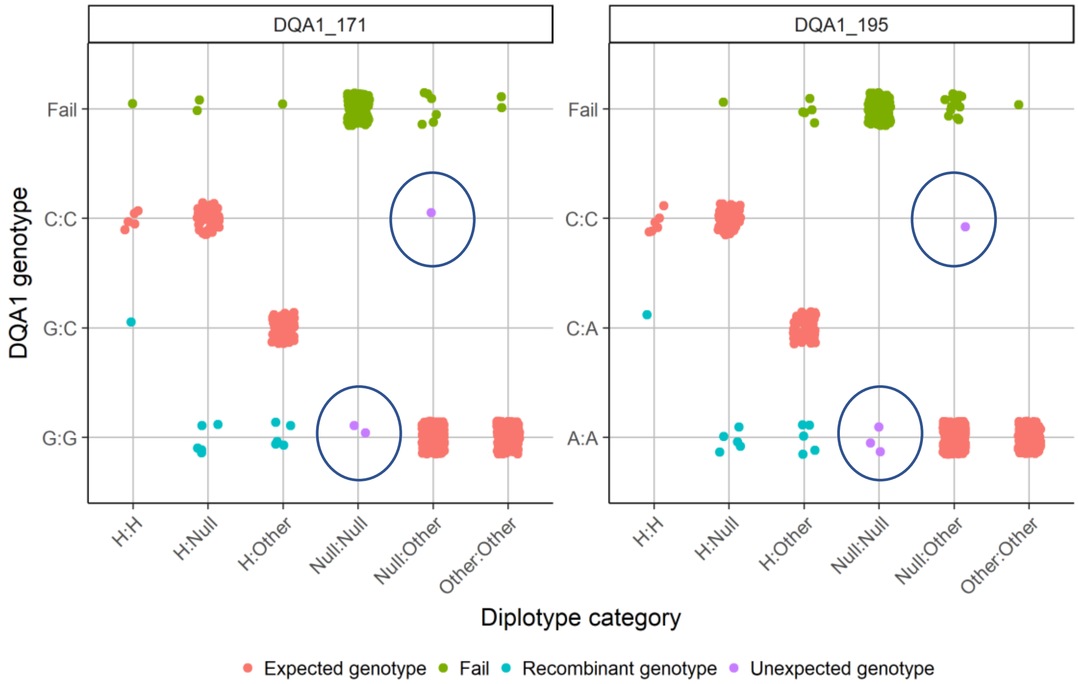


Figure S5. Comparison of the SNP diplotype category with DQA1 genotypes to detect recombinant B to H SNP haplotypes. Haplotypes are grouped as H, which is involved in the recombination event, or according to whether or not they carry a DQA1 locus (Other includes haplotypes B, C, D and F, and Null includes A, E and G). Five individuals had unexpected DQA1 genotypes (circled), either Null:Null or Null:Other haplotype combinations. Points are shown jittered to indicate samples sizes.
